# Attention expedites target selection by prioritizing the neural processing of distractor features

**DOI:** 10.1101/2020.07.16.206573

**Authors:** Mandy V. Bartsch, Christian Merkel, Mircea A. Schoenfeld, Jens-Max Hopf

**Affiliations:** Leibniz Institute for Neurobiology, 39118 Magdeburg, Germany; Department of Neurology, Otto-von-Guericke University, 39120 Magdeburg, Germany; Kliniken Schmieder Heidelberg, 69117 Heidelberg, Germany

**Keywords:** EEG, attention, feature-based attention, human, neuroscience, visual cortex, color, vision

## Abstract

Whether doing the shopping, or driving the car – to navigate daily life, our brain has to rapidly identify relevant color signals among distracting ones. Despite a wealth of research, how color attention is dynamically adjusted is little understood. Previous studies suggest that the speed of feature attention depends on the time it takes to enhance the neural gain of cortical units tuned to the attended feature. To test this idea, we had human participants switch their attention on the fly between unpredicted target color alternatives, while recording the electroencephalographic brain response to probes matching the target, a non-target, or a distracting alternative target color. Paradoxically, we observed a temporally prioritized processing of distractor colors. A larger neural gain for the distractor followed by stronger attenuation expedited target identification. Our results suggest that dynamic adjustments of feature attention involve the temporally prioritized processing and elimination of distracting feature representations.

## Introduction

When searching for tomatoes in a crowded veggie counter, one will most likely rely on their red color to spot them. That is because our brain can easily select a specific color amongst competing distractor color signals (green cucumbers, orange carrots, etc.), and guide our attention to locations of its occurrence (J. M. Wolfe & Horowitz, 2004; Jeremy M. Wolfe, 1994). Such guidance by color is particularly efficient, since attention to color can operate in parallel across the entire visual field irrespective of item locations, thereby allowing for a rapid localization of colored objects. The location-independent nature of such feature selection processes is referred to as spatially global feature-based attention (GFBA), and its underlying neural correlates have been well-characterized both in the human (Andersen et al., 2013, 2015; Bartsch et al., 2015; Bondarenko et al., 2012; Hopf et al., 2004; Peelen et al., 2009; Saenz et al., 2002; Serences & Boynton, 2007; White & Carrasco, 2011), and the monkey (Bichot et al., 2005; Martinez-Trujillo & Treue, 2004; Maunsell & Treue, 2006; C. J. McAdams & Maunsell, 2000; Carrie J. McAdams & Maunsell, 1999; Treue & Martínez Trujillo, 1999). At the single neuron level, GFBA is assumed to arise from a multiplicative gain enhancement of feature selective units in visual cortex tuned to the attended feature value (Martinez-Trujillo & Treue, 2004; C. J. McAdams & Maunsell, 2000; Treue & Martínez Trujillo, 1999). Consistently, EEG/MEG experiments in humans revealed that GFBA is associated with gain enhancements of the neural population response for the attended feature, starting around 150-200ms after stimulation onset in extrastriate visual cortex areas (Bartsch et al., 2015, 2017, 2018; Bondarenko et al., 2012; Garcia-Lazaro et al., 2016; Hopf et al., 2015). It is reasonable to assume that efficient target identification will depend on how rapidly this global biasing can be build up for the attended feature. Still, the processes that serve to adjust feature selectivity dynamically among multiple attended feature values are little understood.

In most GFBA experiments, the target is defined by a single constant feature value (e.g., one color) for a complete experimental trial-block, such that participants implement a stable preset-bias for that specific target feature value. In everyday life, however, target items appear in different shades, and we often look for several things simultaneously. For example, with both red tomatoes and green avocados being on our shopping list, we often don’t know in which order we will find those items. As a consequence, we are required to hold a parallel bias for both colors (attentional template for red and green), but to adapt (bias) color selectivity ‘on the fly’ to the color of the vegetable that happens to be encountered in a given moment. That is, when first coming across tomatoes, our brain should strengthen the color bias for red, while attenuating the response to green, a currently-distracting target color alternative (see Figure 1). But how does the brain quickly shift color selectivity between colors when not knowing before which of them will be target-defining and which will be distracting?

**Figure 1.**
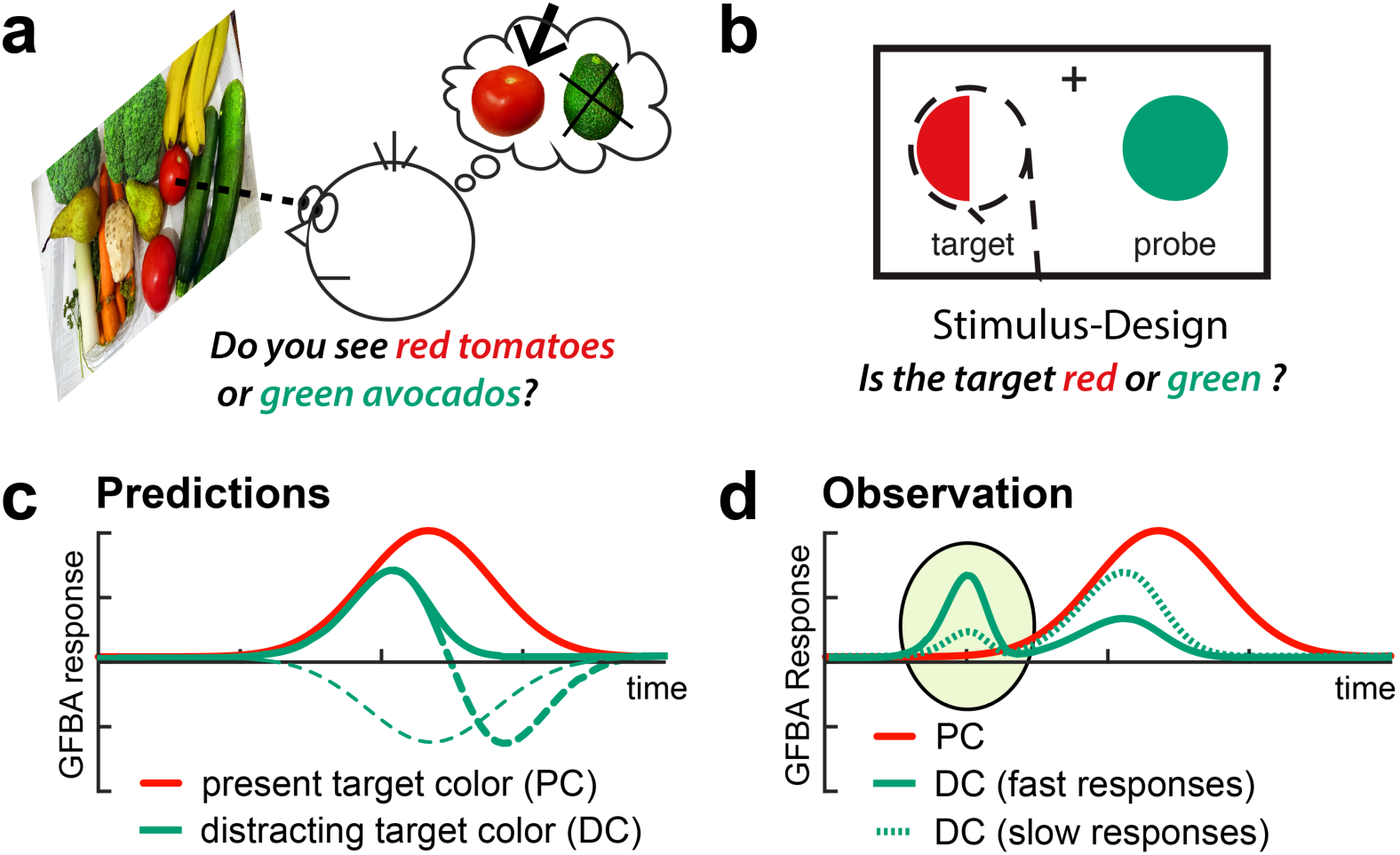
Dynamic of attentional color biasing. **(a) Motivation**. When searching for both red and green items, not knowing what we will encounter first, our brain must decide ‘on the fly’ which color is currently contained in a target object (here: red) and which color would be rather distracting (here: green), and adjust the color bias in the brain accordingly. **(b) Experimental idea**. To investigate this color biasing dynamic independent of other influences like object location, we created simplified stimuli where the target location was fixed, but its color changed unpredictably between two colors (see Figure 2 for details). **(c) Predictions**. The color selection bias in the brain was assessed as the amplitude of the global feature-based attention (GFBA) response to that color (for details see Materials and Methods). Participants may initially bias both possible target colors (here: red and green). The response to the distracting color alternative (DC, here: green), will then decay (green solid) as the neural bias for this color declines in favor of the present target color (PC). Alternatively, the DC might become actively suppressed below baseline either with a delay (green thick dashed), or right from the beginning of the GFBA modulation (green thin dashed). **(d) Observation**. Contrary to our predictions, the processing of the distracting target color alternative (DC, green) gained temporal priority (marked by the ellipse). On trials with a fast response time -i.e., fast identification of the target’s color-, participants showed a prominent early selection of the DC followed by its stronger attenuation in the time range of maximal biasing of the present target color (PC, red). For slow responses (green dashed), the early response to the DC and its subsequent attenuation were less pronounced.

The aim of the present study is therefore to investigate the cortical dynamics of adjusting color selectivity among two target colors in the moment one or the other color has to be selected. To accomplish this, we employ a modified version of the unattended probe paradigm (UPP) used in Bartsch et al. (2015, experiment 3), with the task requiring a color discrimination among two target color-values. We informed participants that the upcoming target will be drawn in one of two possible target colors (e.g., red or green). A simultaneously presented, unattended color probe could be drawn in either of those colors. Analyzing the brain response to the probe allowed us to assess the temporal evolution of the GFBA response as a function of whether the color must be biased for target identification (probe matches present target color, PC), or de-emphasized in favor of the presented target color (probe contains distracting target color alternative, DC) on a given trial. We also add a control condition, where subjects view exactly the same stimuli, but are asked to discriminate the shape of the target, while color is completely task-irrelevant. On each trial, there is a 50% chance that one or the other target color appears. Hence, it would be a reasonable assumption that participants implement a balanced top-down bias (attentional template) for both target colors. In the moment the target appears, however, the bias must shift to the color-value of the currently-presented target. Regarding the time course and amplitude of the GFBA response of the two colors, several scenarios are possible, which are illustrated in Figure 1c. There may be an initial GFBA response for both the present (PC, red) and the distracting (DC, green) target color appearing with the same temporal onset. The DC, however, may raise to a smaller amplitude and soon fade away as the neural bias for this color declines in favor of the PC (green solid), fitting previous observations in Bartsch et al. (2015). A related possibility is that the DC does not only fade but will be actively suppressed below the level of an unbiased color (green thick dashed). As an extreme, this suppression of the DC could already start at the onset of the GFBA response (green thin dashed).

## Results

### Dynamics of color attention during color and orientation discrimination

Twenty-two human adult volunteers participated in both the color and the orientation task (see Figure 2 for experimental design). They were required to attend to a semi-circle presented in the left visual field (VF) and to either identify its color (color task), or orientation (orientation task). Importantly, the target was always randomly drawn in one of two possible target color alternatives (e.g., red or green), such that after stimulus onset, participants had to quickly adapt their bias ‘on the fly’ toward the present target color (PC), and away from the distracting target color alternative (DC). The temporal development of color selectivity was tracked by recording the electroencephalographic response to simultaneously presented irrelevant color probes (unattended probe paradigm, UPP, for details see Materials and Methods). This resulted in three different trial types per experimental condition, i.e., the probe could contain the PC, the DC, or a non-target color. The response difference measured between attended colors (PC and DC) and irrelevant non-target colors was taken as global feature-based attention effect (GFBA) and served to track cortical color biasing. When participants performed the color task, we expected to observe a dynamical adjustment of attentional color selectivity in favor of the PC (see Figure 1c). The orientation task served as an experimental control condition rendering the color of the current target irrelevant. That way, we could distinguish between effects related to color attention (color task) and effects that might arise when discriminating the stimuli without explicitly attending to their color (orientation task).

**Figure 2.**
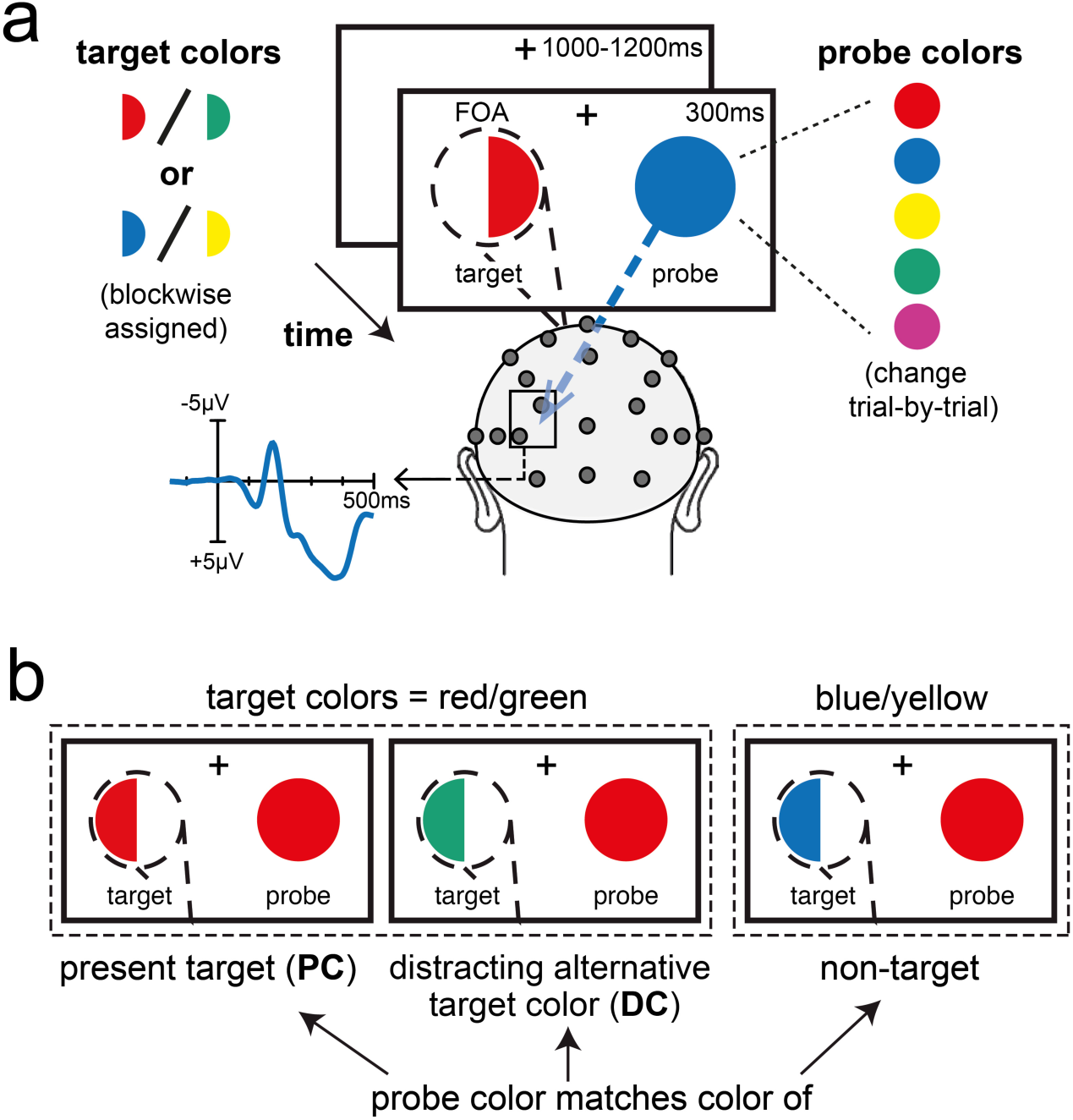
Experimental design. **(a)** Participants attended to a colored hemi-circle presented in the left VF (dashed line = spatial focus of attention, FOA) and reported by button press either its color (color task blocks) or the side (left/right) of its convexity (orientation task blocks). The target varied trial-by-trial unpredictably between the two blockwise-assigned target colors (i.e., between red and green, or between blue and yellow). On each trial, the color probe simultaneously presented in the right VF was randomly drawn from five colors (red, green, blue, yellow, magenta). Effects of global feature-based attention (GFBA) – as a measure for attentional color selectivity in visual cortex – were assessed by comparing the event-related brain response elicited by an unattended color probe as a function of whether it matched the present target (PC), the distracting alternative target color (DC), or neither of them (non-target). **(b)** Trial types. The probe (here: red) could either contain the PC, the DC (here: red probe but green target in an attend red/green block), or could represent a non-target color currently not relevant (here: red probe in an attend blue/yellow block).

### Behavioral results – distracting color alternative impairs performance in the color attention task

Figure 3 displays response time (RT) and response accuracy for the different trial types. As can be seen, participants responded fast (<410ms) and with high accuracy (>92% correct) across all conditions. However, responses seemed to be slightly slower and less accurate when performing the color task, which was most obvious when probes contained the distracting target color alternative (DC, grey bars). Conducting 2×3 rANOVAs with the factors TASK (orientation/color) and COLOR (PC/DC/non-target) revealed significant main effects for TASK (accuracy: F[1,21]=5.15, p=0.034, RT: F[1,21]=11.17, p=0.003), confirming the performance decrement in the color task. As expected, there was also a main effect of COLOR (accuracy: F[2,42]=5.08, p=0.011; RT: F[2,42]=7.56, p=0.002) that showed a significant interaction with TASK for response time (F[2,42]=9.44, p=0.002) but not response accuracy (F[2,42]=2.60, p=0.101). Subsequent t-tests confirmed that, for the color task, responses were significantly slower and less accurate when probes matched the DC compared to when probes matched the PC (accuracy: p = 0.002; RT: p = 0.001), or a non-target color (accuracy: p=0.049; RT: 0.009) with no difference between the latter (accuracy: p = 0.108; RT: p = 0.056). For the orientation task, we did not observe significant influences of the probe color on performance besides a slight response time increase when the probe contained the PC compared to a non-target color (p = 0.016, all other p > 0.17). That is, when asked to discriminate the target’s shape but not its color, a probe matching its color might have been slightly distracting.

**Figure 3.**
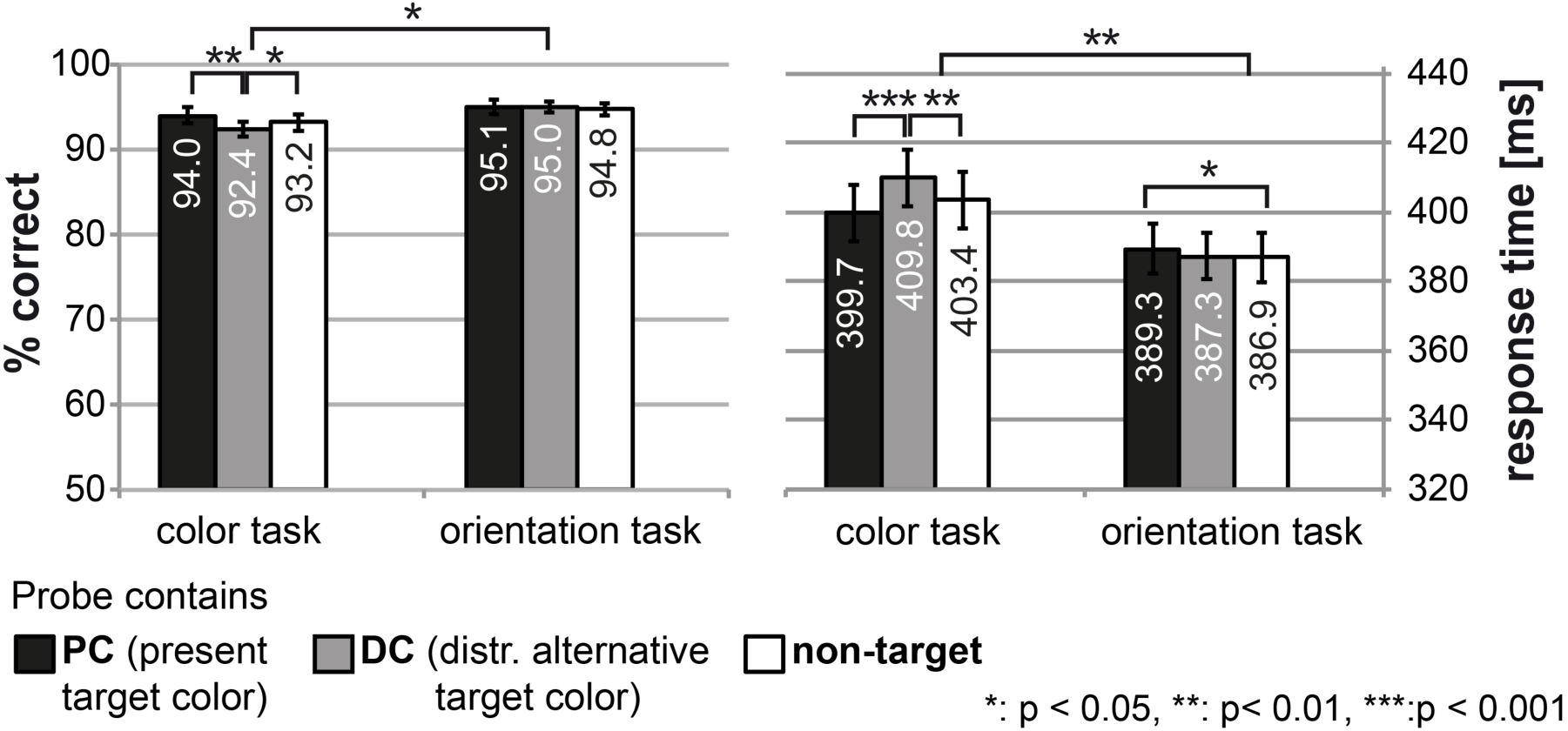
Behavioral performance. Shown are the percentage of correct responses (left) and response times (right) of both the color and the orientation task for all trial types. Participants responded highly accurately and fast across all conditions. However, the performance was slightly lower in the color task, most prominent as a response delay on trials where the probe matched the distracting alternative target color (DC, grey bars). The error bars represent the standard error of the mean (SEM).

Taken together, though behavioral performance was very high in both tasks, when participants performed the color task, the DC significantly slowed performance, indicating that presenting the target color alternative did, indeed, distract the processing of the present target color, which slightly delayed responses. Consistently, no such DC performance decrement was observed under conditions of the orientation task, indicating that participants successfully implemented different attentional task sets with the alternative color of the target being irrelevant and, hence, not distracting during attention to orientation.

### Event-related potential responses (ERPs)

According to previous work (Bartsch et al., 2015, 2017, 2018; Bondarenko et al., 2012), we expected to find a negative voltage enhancement for probes containing attended compared to unattended colors at parieto-occipital electrode sites contralateral to the probe location (here: signal averaged across PO3 and PO7). Specifically, when subjects performed the color task, we should observe modulations for both target color alternatives (PC and DC are both part of the attentional set). In contrast, when discriminating the orientation of the target irrespective of its color, color biasing should, if at all, only be present for the PC (i.e., for the color contained in the object under discrimination). An overall sliding window 2×3 ANOVA (see Materials and Methods) with the factors TASK (color/orientation) and COLOR (PC/DC/non-target) revealed a significant early TASKxCOLOR interaction (73-96ms) and a late main effect for COLOR (167-254ms). Subsequent analyses focusing on those time ranges revealed pronounced early and late processes of color biasing for the color task, and a more general task-independent enhancement of the PC in the later time range, as detailed in the following.

### Color task – early cortical bias for the distracting color

Figure 4a shows the ERP responses for the color task. When participants had to decide which of two colors was present in the focus of attention, there was first a negative enhancement of the distracting color alternative (DC, grey line) in the early time range (73-96ms), followed by a negative enhancement most prominent for the present target color (PC, black solid line) in the late time range (167-254ms). The brain response to unattended non-target colors (black dashed line) served as reference. To isolate the GFBA effects, the response to the unattended non-target color was subtracted from that of the DC (Figure 4b, upper row) and the PC (Figure 4b, lower row). As can be seen in the resulting difference waveforms, the early color bias was only present for the DC (p=0.0179) but not for the PC (p=0.6521). The late bias, however, was apparent for both DC (p=0.0403) and PC (p<0.0005), albeit much more pronounced in the latter (mean amplitudes differ significantly, p=0.0103). The respective topographic field distributions of the GFBA responses reveal similar field distributions for early and late enhancements with a parieto-occipital maximum at electrodes PO3 and PO7 (Figure 4b, on the right). Although there seems to be a small early modulation for the PC outside the time range of investigation (Figure 4b, lower row), an additional explorative sliding window t-test (sample-by-sample, 11.8ms window) between 0-130ms revealed no significant modulation before or shortly after the early time window (all p>0.05).

**Figure 4.**
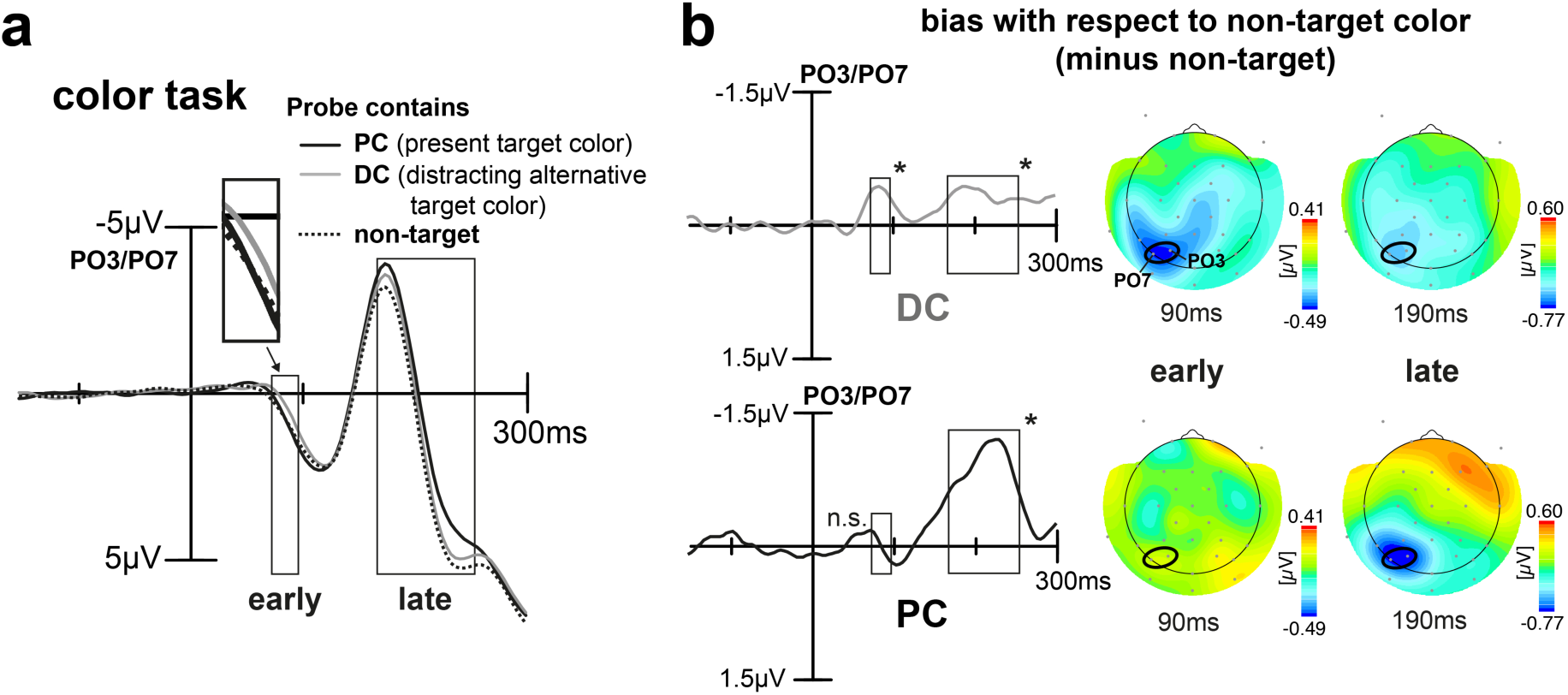
ERP results for the color task. **(a)** Shown is the ERP elicited by the probe at PO3/PO7 (signal averaged) for the different trial types when participants were to discriminate the color of the target. Rectangles highlight time ranges of significant brain response variations as derived by the 2×3 rANOVA. Surprisingly, participants show a pretty early bias (higher negativity around 73-96ms) for the DC (grey line), see inset for an amplified depiction. Significant biasing for the color of the PC emerges later (167-254ms). **(b)** Difference waveforms for DC minus non-target color (upper row) and PC minus non-target color (lower row). The respective topographical field maps on the right display representative time points at early and late modulation maxima, positions of electrodes used for analyses are highlighted (black ellipses). In the early time range, there is a prominent bias for the DC but not the PC. In the late time range, this pattern becomes inverted with a strong selection bias for the PC and only a weak modulation for the DC.

The modulation observed in the N1/N2 time range (here referred to as “late” effect) is well in line with previous literature, where such negative enhancements have been observed for the attended color (Bartsch et al., 2015, 2017, 2018). Also fitting previous observations, this effect is smaller and more transient for a color that is part of the attentional set but not discriminated on a given trial (here: DC) (cf. Bartsch et al., 2015, experiment 3). The response in the N1/N2 time range therefore matches the prediction illustrated in Figure 1c (red and green solid, cf. with Figure 5a late time range), with an initial rise of the response for both attended colors, but a smaller amplitude for the DC that is, however, never suppressed below the level of non-target color responses. To our surprise, and not fitting any of our predictions, those late modulations were preceded by an early negativity around 70-100ms, which appeared only for the DC but not the PC. This seems to be counterintuitive as it suggests that the DC gains an early selection bias above the PC. Given the 50% chance that the one or the other target color appears on a given trial, one would expect that the top-down bias for both colors is overall balanced, and that upon stimulus onset, the neural processes mediating the selection of the PC are involved as fast as possible. The here observed response pattern, instead, suggests that the DC undergoes prioritized processing. One possibility would be that the DC is selected with temporal priority in order to rapidly build a representation that serves its rejection from further processing (selection for rejection), which would ultimately facilitate the selection of the PC. In fact, such temporally prioritized selection of distracting information was previously observed in visual search (Donohue et al., 2018), where a stronger early distractor selection was associated with faster target discrimination. A similar function could underlie the here observed early selection bias of the DC. If this is the case, the amplitude of the early negativity should inversely relate to the time it takes to discriminate the PC. Furthermore, if the early negativity reflects the selection for rejection of the DC from current processing, a higher the amplitude of the early negativity should be associated with a smaller later GFBA response to this color. To address this possibility, we compared the GFBA response to the PC and DC after separating the data into fast and slow correct responses (Median response time split analysis, see Materials and Methods). Figure 5 shows respective brain responses for the color task of trials where subjects gave fast versus slow responses.

**Figure 5.**
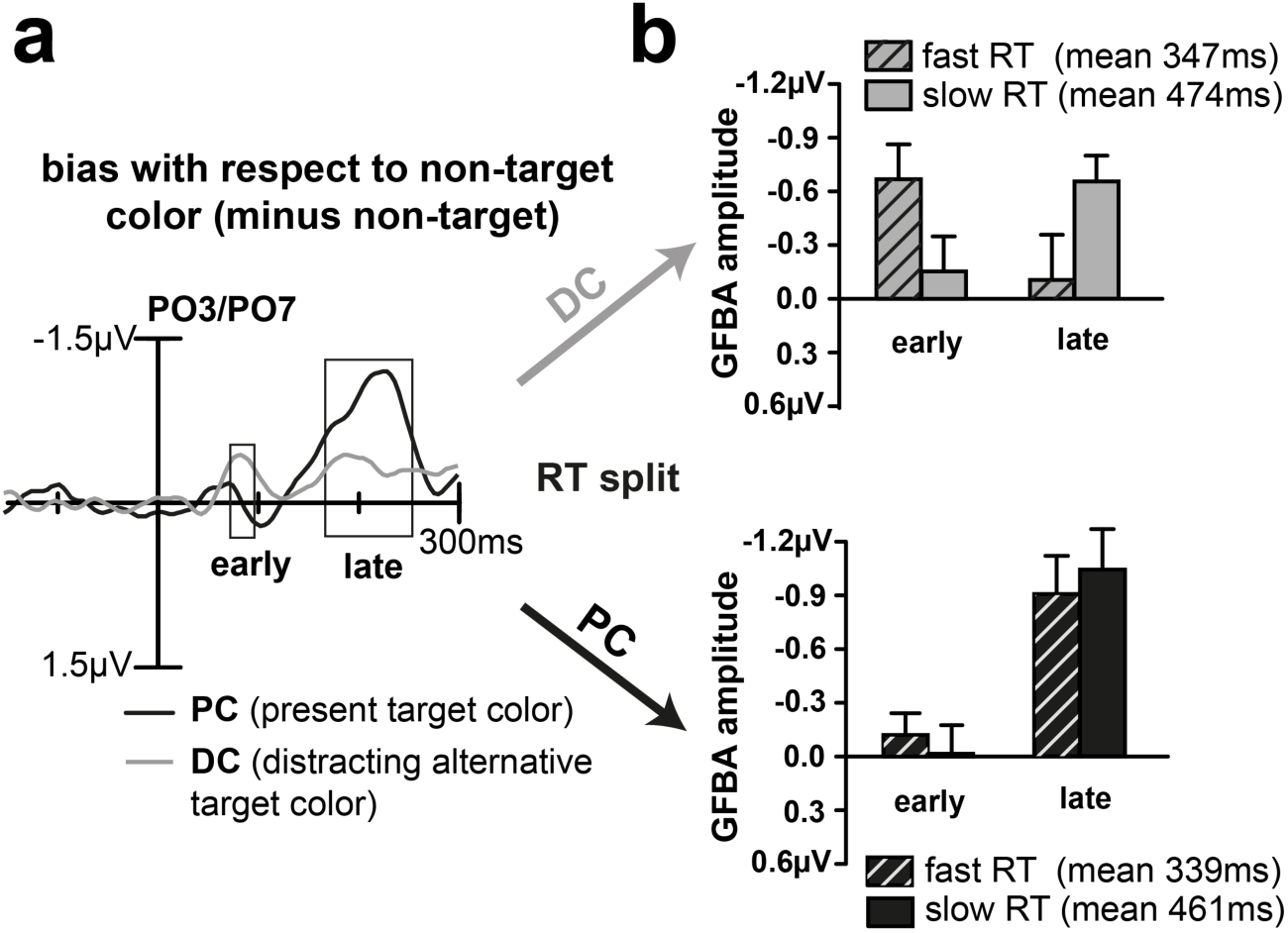
Median response time split analysis. (a) Difference waveforms for the PC (black) and DC (grey) replotted together from Figure 4b for better comparison. Rectangles indicate previously determined early and late time ranges of significant experimental variation. (b) Median split into fast and slow response times (RT) for DC (grey, upper row) and PC (black, lower row). Mean GFBA amplitudes of the early and late time range are shown for fast (striped) and slow (plain) responses. For DC, the early negative enhancement was higher when participants responded fast, which was inversely correlated to the size of the late bias (significant early/late GFBA amplitude x fast/slow RT interaction, p = 0.00035, see text for details). For the PC, in contrast, there was no significant difference in GFBA amplitudes between fast and slow responses.

### Strong early distractor color biasing and subsequent attenuation expedites target color identification

As can be seen in Figure 5b, upper row, for the DC, the mean amplitude of the early selection bias did, indeed, vary with response time as predicted. On fast responses (striped bars), the early modulation was significantly higher (p=0.0132), while the late selection was reduced relative to slow responses (marginally significant, p=0.0558). A rANOVA on ERP amplitudes with the main effects EARLYLATE (early GFBA amplitude, late GFBA amplitude) and FASTSLOW (fast responses, slow responses) confirmed a significant EARLYLATExFASTSLOW interaction (F[1,21]=18.15, p = 0.00035) with no significant main effects (EARLYLATE: F[1,21]=0.164, p = 0.689; FASTSLOW: F[1,21]=0.001, p = 0.982) for the DC. For the PC, on the other hand, there was no significant early modulation at all but a strong late bias that did not vary with response time. As expected, the analogous rANOVA found only a significant main effect for EARLYLATE (F[1,21]=33.13, p < 0.0005) but not for FASTSLOW (F[1,21]=0.004, p = 0.953), or the interaction (F[1,21]=0.37, p = 0.547). Importantly, with a mean fast response time of 347ms on DC trials, responses were initiated well after (and not before) the early modulation, probably during or shortly after the late time range, in line with an influence of the early cortical distractor processing on the speed of target identification.

Together, the RT split pattern clearly supports the idea that a preferential biasing of the DC and its subsequent rejection expedites the identification of the target’s color. This conclusion implies that it is the prominence of the early sensory representation of the DC and not so much the very processing of the PC itself that is important for efficient target identification. If, however, the early highlighting of the neural DC representation is crucial for task performance, this leads to a notable prediction: When one would strengthen the cortical representation of the PC, instantiating the temporary priority of the DC would require to counteract this stronger PC bias, and hence, lead to a larger early modulation for the DC. If, however, the early distractor prioritization is not mandatory for task performance, a stronger representation of the target might as well eliminate the need for the early DC biasing. One way to manipulate the strength of target representation would be to exploit the effects of inter-trial color priming (Kristjánsson, 2006; Nakayama & Mackeben, 1989; Pinto et al., 2005; Theeuwes, 2013). Specifically, repeating the target feature (PC) should strengthen its sensory representation, leading to faster response times and decreased distractor interference.

To determine the influence of color priming on our modulation sequence, we split conditions into trials where the target was repeated and trials where the target color switched (Figure 6a, for details see Materials and Methods, Target repetition analysis). Participants responded about 40ms faster on target repetition trials, irrespective of probe color (PC: 381ms vs. 419ms, DC: 390ms vs. 428ms, non-target: 387ms vs. 419ms). A two-way rANOVA with the factors COLOR (probe matches PC/DC/non-target) and REPETITION (target repeated/switched) revealed a significant main effect of REPETITION (F[1,21]=43.81, p<0.0005) and of COLOR (F[2,42]=9.32, p=0.001), but no interaction between them (F[2,42]=1.02, p=0.351). As visible in Figure 6b, repeating the target color increased the early selection and subsequent attenuation of the distracting color. Hence, it seems that in fact the stronger neural representation of the PC on primed trials does not abolish the early DC selection but rather enforces the DC biasing pattern, underscoring its role for efficient target selection.

**Figure 6.**
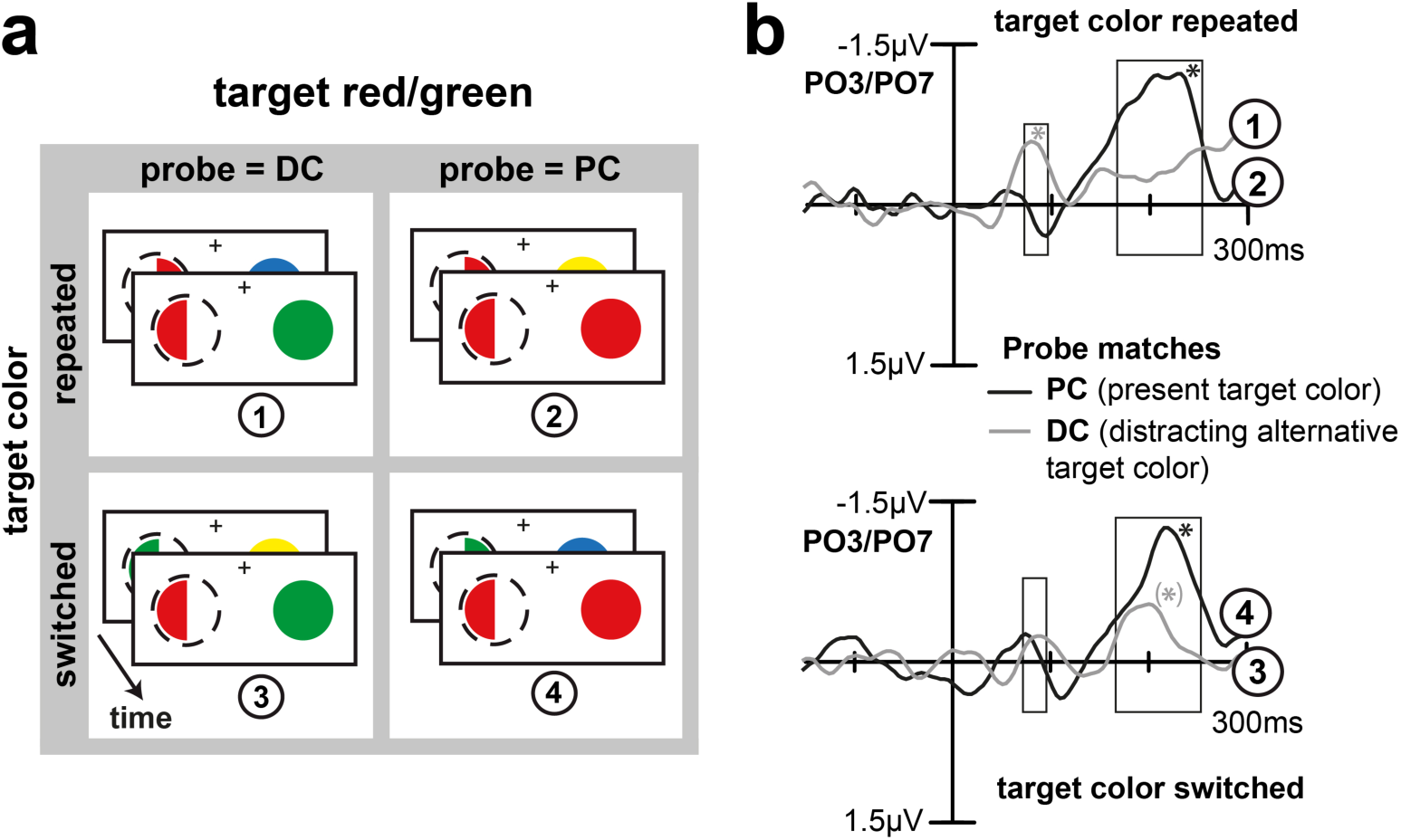
Target repetition analysis. **(a)** Trials in which the probe contained the DC, or the PC were sorted according to whether the color of the previous target was repeated (stronger color bias for the PC), or switched (weaker color bias for the PC). Here exemplarily shown for trials of an attend red/green block and a red target. **(b)** Difference waveforms (minus non-target) for PC (black) and DC (grey) probes for repeat (upper row) and switch trials (lower row). Rectangles indicate previously determined time ranges of early and late color biasing. A stronger bias for the PC entailed a pronounced early color biasing for the DC (p = 0.0130), much smaller and not statistically significant on switch trials (p = 0.2761). Corroborating the response time split (see Figure 5), a strong early selection of the DC was followed by its weaker selection in the late GFBA time range and linked to faster target identification. Stars indicate significant mean amplitude modulations in the indicated time ranges (p<0.05), the late GFBA response for DC under switch conditions (lower row, grey line) was only significant when considering a shorter time window (i.e., from 167-210ms).

Taken together, the ERP responses observed during the color task indicate that a strong initial selection of the DC entails its weaker late biasing/less interference of this distracting color, and leads to faster response times (Figure 1d). The pattern is suggestive of a mechanism akin to “selection for rejection” previously observed for distracting color singletons in visual search displays (Donohue et al., 2018).

### Orientation task - rendering target color irrelevant abolishes early color biasing effects

To draw strong conclusions, it is important to verify that the GFBA effects observed in the color task, indeed, reflect pure modulations of attention to color and are not based on low-level sensory processes that differ among the experimental conditions. In particular, trials with the probe matching the PC versus the DC differ as to the diversity of color in the stimulus array. On PC trials, the same color appears in the left and right VF, while on DC trials different colors appear (see Figure 2b). It is therefore critical to rule out that differences in the early modulation time range are triggered by such color imbalance between VFs. To this end, we analyzed the brain responses to the very same physical stimuli when subjects were asked to discriminate the orientation of the target item, while color was completely task irrelevant. Here, we expected no enhancement for the alternative target color (DC). However, as observed in previous work (Bartsch et al., 2015), we anticipated some later GFBA effect when probing the color of the target under discrimination (PC). Figure 7a shows brain responses averaged over trials with the same color assignment as in the color task, but when participants performed the orientation discrimination with color being irrelevant. As can be seen, there was no obvious early color bias, especially no negative enhancement of the DC (grey line). Instead, the difference waveforms in Figure 7b reveal a rather small counter-modulation (positivity) in the early time range for both PC and DC which is, however, only significant for the DC (p=0.0242). As expected, there was some late bias for the PC in the late N1/N2 time range (black solid line), but no effect for the DC (late time window, grey line). The late PC enhancement is, however, smaller compared to that of the color task (cf. Figure 4b, lower row), which is confirmed by a paired t-test (mean amplitudes differ, p=0.0031). The late enhancement in the orientation task is preceded by a smaller negative enhancement around 130ms. An explorative sliding window t-test in the time range between the early and late time window (sample-by-sample, 11.8ms window), however, failed to reach significance. Importantly, the DC biasing pattern (early selection and subsequent attenuation) is eliminated when participants did not attend to color and thus highly unlikely to be caused by mere low-level sensory differences among trial types.

**Figure 7.**
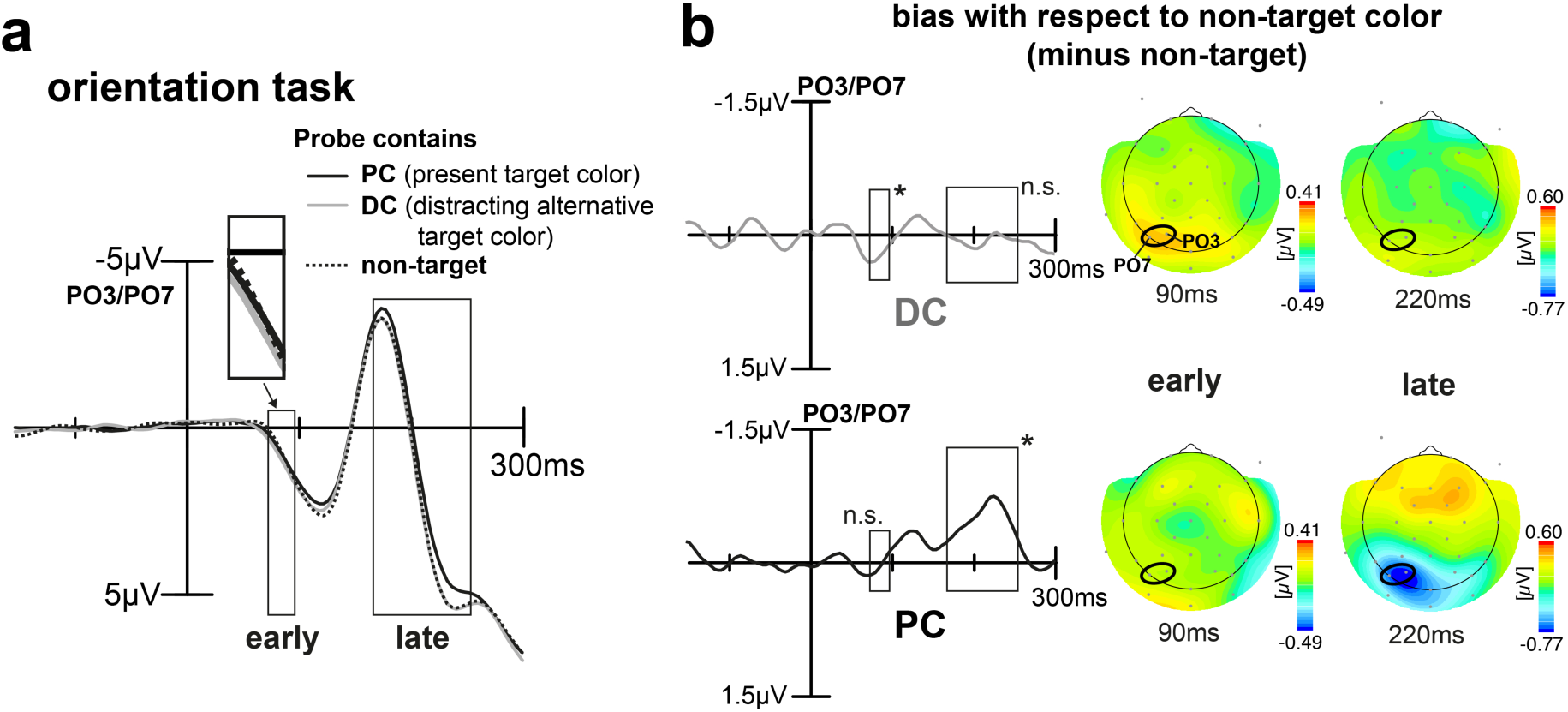
ERP results for the orientation task. **(a)** Shown is the ERP elicited by the probe at PO3/PO7 (signal averaged) for the different trial types when participants were to discriminate the orientation of the target. Rectangles highlight time ranges of significant brain response variations as previously derived by the 2×3 rANOVA. This time, participants show no negative enhancement of the DC (grey line), see inset for an amplified depiction. Again, there is a late negative enhancement for the color of the present target object (PC) (167-254ms). **(b)** Difference waveforms for DC minus non-target (upper row) and PC minus non-target (lower row). The respective topographical field maps on the right display representative time points at early and late modulation maxima with the positions of the electrodes used for the analyses being highlighted (black ellipses). The difference waveforms reveal some sort of general small anti-bias (positive modulation) for color in the early time range and a late enhancement that is present for the color of the object being under discrimination only.

To sum up, when participants have to decide among two possible target color alternatives (color task), they appear to first implement a very early bias (73-96ms) for the potentially distracting target color alternative (DC) entailing a reduced late biasing for that color in the N1/N2 time range (167-254ms), where previous work reported template matching and feature discrimination processes to occur (cf. Bartsch et al., 2015, 2017; Bondarenko et al., 2012). Contrary to any of our predictions, the bias for the color currently present in the target object (PC) is built up more slowly resulting in a strong late selection (167-254ms) that would also be present-though smaller in amplitude-when performing a mere orientation discrimination of the object.

## 3. DISCUSSION

The aim of this study was to investigate the temporal dynamics of biasing color selectivity in favor of the currently-presented target color, when an upcoming target appeared unpredictably in one out of two possible colors. To this end, we compared the event-related brain response (ERP) to spatially unattended probes drawn in the target color alternatives, when participant had to decide ‘on the fly’ on a given trial which of them was currently contained in the target (present target’s color, PC), and which was rendered a now distracting color alternative (DC). We expected observers to establish an initially balanced top-down bias for both colors, with the response to the DC soon fading away or being even actively suppressed (Figure 1c). ERP responses showed, indeed, that observers established a top-down attentional bias for both colors, which is in line with our previous observations (Bartsch et al., 2015; Bondarenko et al., 2012) as well as work showing that observers can establish attentional control settings for multiple colors at once (Adamo et al., 2008; Grubert & Eimer, 2016; Irons et al., 2012; Moore & Weissman, 2011; Ort et al., 2019), and that guidance by two target colors is possible in visual search (Beck et al., 2012; Stroud et al., 2012). In the present study, both the PC and the DC, elicited global feature-based attention (GFBA) modulations roughly at the same latency in the N1/N2 time range (167-254ms, here referred to as late effect). In line with Bartsch et al. (2015, experiment 3), the response to the DC was smaller in amplitude and decayed faster (Figure 5a, late time range), fitting the prediction depicted in Figure 1c, solid lines.

However, beyond our predictions, we observed that the DC but not the PC elicited a prominent and rather early modulation (peaking around 90ms) roughly ∼80 ms prior to the onset of the GFBA effects in the N1/N2 time range (Figure 4b, upper row). The polarity and scalp topography of this early modulation was very similar to that of the subsequent GFBA effects, suggesting that it reflects some form of early global color selection which is functionally related to the following GFBA response. At first glance, this observation is counterintuitive, as one would expect that the prioritized selection of the DC impedes and delays the discrimination of the PC. However, further analyses revealed the opposite effect - a stronger early DC bias was associated with a faster target identification. What then, is the function of this early modulation? One possible explanation would be that the priority processing of the distracting target color alternative serves to build a temporary representation for eliminating or attenuating the selection bias for this color (selection for rejection). This would reasonably be most effective before the ‘regular’ cortical processes (N1/N2 GFBA effects) dealing with the representation and discrimination of the color of the present target start to arise. A blocking of DC processing in the GFBA time range would effectively bias the competition towards the PC (Desimone & Duncan, 1995; Kastner & Ungerleider, 2001), thereby expediting target discrimination. The median RT-split analysis of the present data supports this interpretation. A stronger early modulation in fast RT trials was associated with a smaller later GFBA amplitude (see Figure 5b, upper row, and Figure 1d for a schematic depiction), suggesting that the GFBA response to the DC was, indeed, attenuated in the N1/N2 time range. Importantly, for the PC, no significant variation of the GFBA response as a function of RT was seen, indicating that the processing of the DC but not that of the PC was linked to response time differences. Hence, the present data suggest that selecting among unpredictably changing target colors with equal top-down bias, involves a temporarily prioritized representation of the currently irrelevant/distracting color. This early representation presumably serves to suppress later GFBA responses to this color, thereby ultimately facilitating the discrimination of the actual target color.

The analysis of trial-by-trial color priming in the present experiment supports this interpretation. If the early bias reflects, indeed, an active process to highlight the distracting target color alternative, it should be influenced by the strength of the cortical representation of the present target color. A priming-driven bias toward the PC on repeat trials (Kristjánsson, 2006; Nakayama & Mackeben, 1989; Theeuwes, 2013) led to a stronger early DC modulation (and subsequent attenuation) consistent with activity counteracting the enhanced sensory bias for the PC on repeat trials to instantiate the priority representation of the DC. Importantly, the early DC selection was not itself a mere artifact of DC color priming, since the DC was not systematically repeated on those trials in the unattended color probe (previous probe color random, cf. Figure 6a).

It is worth noting that such temporal prioritized selection of distracting items has been reported in visual search (Donohue et al., 2018). A salient distractor (color singleton) was found to elicit an N2pc-like early (around ∼150 ms) component (N1pc) with an onset prior to the same component elicited by the target singleton. Notably, a bigger N1pc response to the distractor singleton was associated with an earlier Pd (distractor positivity) reflecting faster distractor suppression (Gaspar & McDonald, 2014; Hickey et al., 2009; Hilimire et al., 2012; Jannati et al., 2013; McDonald et al., 2013; Sawaki & Luck, 2010, 2011), and was linked to faster target selection. These observations were discussed in terms of prioritized selection for rejection of the distractor singleton. The early selection of the DC observed here reflects a conceptually similar operation that prioritizes the selection of distracting feature-values for rejection, raising the possibility that prioritized selection for rejection is a generic mechanism underlying the dynamic of attentional selection. Further experimental work is necessary to clarify the relation between the N1pc and the early DC modulation.

The proposed functional role of the early negativity to the DC notably dovetails with the recently proposed signal suppression hypothesis (N. Gaspelin et al., 2017; N. Gaspelin & Luck, 2018; Nicholas Gaspelin & Luck, 2018, 2019; Sawaki & Luck, 2010, 2011), or the related salient-signal suppression account (Gaspar & McDonald, 2014; Jannati et al., 2013). The core idea is that (spatial) attentional capture by irrelevant singletons in visual search can be prevented via active distractor suppression (reflected by the Pd component) before attention would be captured by the item. The present data suggest a conceptually analogous scenario in feature space, with the two target colors being of higher salience in a color-based representation than irrelevant non-target colors. The early highlighting of the DC allows for the attenuation in the time range of the GFBA modulation, whose later portions are known to reflect the discrimination of the target (discrimination matching effect, Bartsch et al., 2015; Bondarenko et al., 2012). A central notion of the signal suppression account is that the distractor suppression must be informed by something that identifies the to-be-suppressed item to invoke top-down attentional control. This has been referred to as ‘attend-to-me signal’ (Jannati et al., 2013; Sawaki & Luck, 2010, 2011) but the neural process underlying this signal is currently unclear. We speculate that the early negativity observed here reflects a neural operation that accomplishes such highlighting operation, conceptually related to the generation of an ‘attend-to-me’ signal in a feature-based representation.

The observation of a small GFBA response in the N1/N2 time range for the DC in the color task replicates analogous observations in Bartsch et al. (2015, eperiment 3) with MEG, using a quite similar experimental setup. In this experiment, however, the DC (there referred to as ‘cross-match color’) elicited no early response modulation in the MEG signal. In Bartsch et al. (2015), EEG data were also recorded, but not reported. A comparison with the present data is therefore worthwhile. Figure 8 plots the GFBA response for the DC and PC of Bartsch et al. (2015) at electrode sites PO3/PO7. Consistent with the present observations, there is a GFBA response starting around 150 ms for both target colors, with the response to the DC being smaller and fading earlier. There is, however, no significant modulatory effect whatsoever before 140 ms after stimulus onset, neither for the PC nor for the DC. Given the similar experimental design, this discrepancy warrants consideration. In Bartsch et al. (2015), participants were to select the shape of the target based on color with shape but not color identity being task-relevant (same response for a red or green target if shape was identical). In contrast, the present color task required subjects to report the color identity of the target. This difference in cognitive operations mirrors a fundamental distinction between selection and (conscious) access as separate attention limits as emphasized by the Boolean map theory of visual attention (L. Huang et al., 2007; Liqiang Huang & Pashler, 2007). According to this theory, conscious access (identification) is bound to a labelled Boolean map (BM) representation of the input, which allows access/identification of only one feature-value per feature-dimension (i.e., one color) at a time. Mere selection, instead, could be based on multiple color values. This would provide spatial and/or shape information about the input, but the identity of the colors would remain inaccessible. That is, when building a BM from attending two target colors (e.g., red and green), shape but not color identity would be accessible, which is sufficient to solve the shape task in Bartsch et al. (2015). For the present color task, in contrast, subjects must access color identity, which requires to build temporally distinct one-by-one maps (Liqiang Huang & Pashler, 2007) for the target colors. We speculate that the early DC modulation reflects such temporally prioritized construction of a BM that identifies the DC for subsequent rejection.

**Figure 8.**
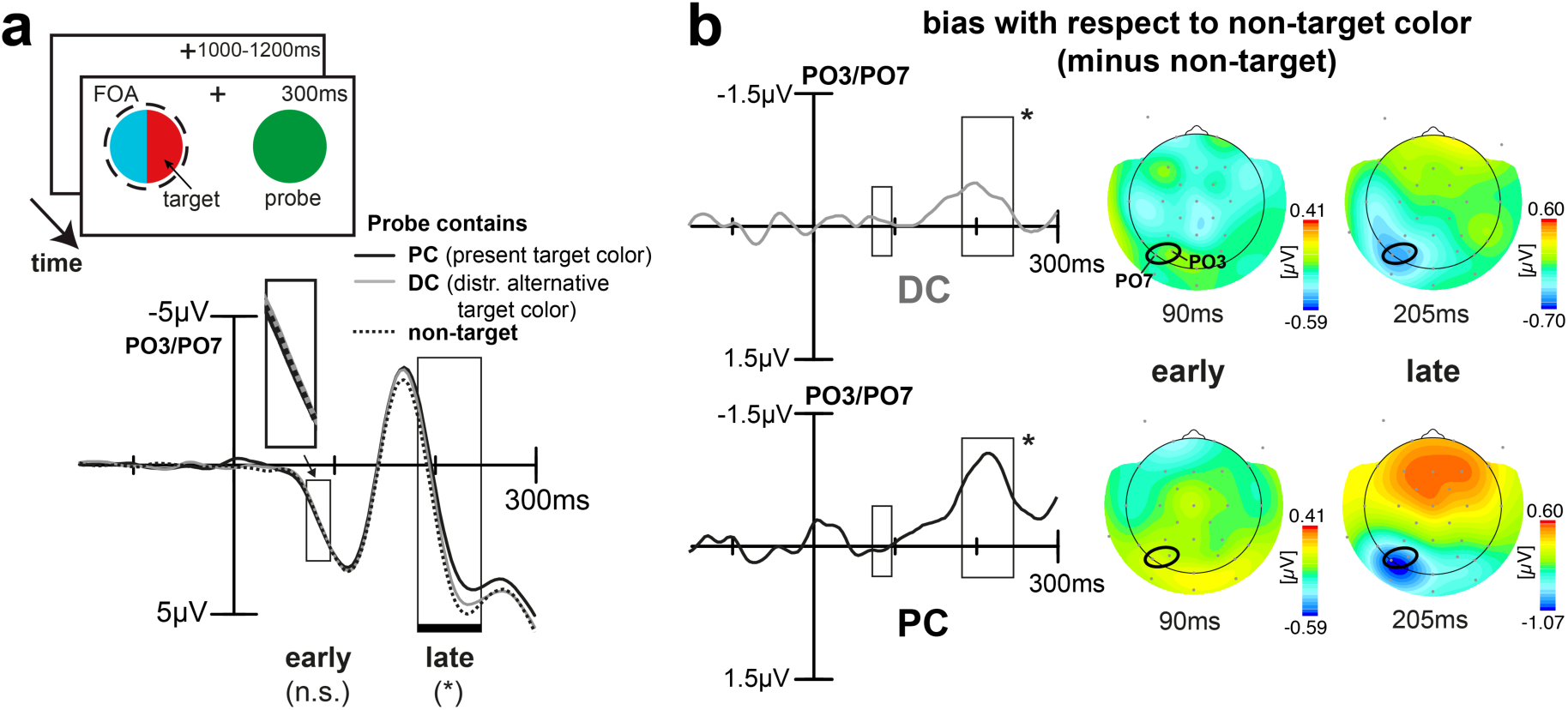
EEG data from Experiment 3 of Bartsch at al. (2015). **(a)** Participants had to report the shape of a color-defined half circle drawn in one of two possible target colors (here: either red or green). The target was combined with a half-circle drawn in a second, task-irrelevant color (here cyan). Shown is the ERP elicited by the probe at PO3/PO7 (signal averaged) for the trials types analogous to the current experiment (PC, DC, or non-target probe). The respective sliding window 1×3 rANOVA (11.8ms window, 0-300ms) revealed only a late significant color-related brain response variation (183-246ms, late rectangle/black horizontal bar, p < 0.02), but none in the early time range as was found in the current experiment (73-96ms, rectangle in early time range). **(b)** Difference waveforms for DC minus non-target color (upper row) and PC minus non-target color (lower row). The respective topographical field maps on the right display representative time points at modulation maxima, positions of electrodes used for analyses are highlighted (black ellipses). The late biasing resembles the current color task with a smaller and faster decaying DC modulation. This time, there is no significant early effect at all.

## Conclusion

Together, our data suggest that trial-by-trial biasing of feature attention ‘on the fly’ among task-relevant color alternatives almost paradoxically elicits a temporally prioritized neural response to the color alternative that is not the target on a given trial. Notably, a large early modulation to the distracting non-target alternative, followed by a reduced GFBA response to it, is found to facilitate target color identification. This suggests that feature competition is resolved by rapidly indexing the distracting color for neural attenuation prior to target color selection which preempts interference in this time period. Such prioritized selection for rejection has also been documented in visual search for distractors and targets competing as salient singletons in the same feature dimension (Donohue et al. 2018). Hence, prioritized selection for rejection may represent a more general attention mechanism called upon when competition within a feature dimension needs resolution.

## Materials and Methods

### Participants

Twenty-two volunteers participated in both color and orientation task (mean age 26.0 years, age range 22-33 years, 12 female, all right-handed). All participants had normal or corrected-to-normal visual acuity and reported normal color vision. Participants gave written informed consent prior to the measurement and were paid 6€/hour for participation (preparation and measurement lasted 2-3 hours). The experimental methods and procedures were approved by the ethics board of the Otto-von-Guericke University of Magdeburg and conducted according to the research regulations of the Declaration of Helsinki. The number of participants as well as the number of trials per experimental condition (> 200) was chosen according to (Luck, 2005), and previous work investigating GFBA components in the EEG/MEG (Bartsch et al., 2015; Bondarenko et al., 2012). Based on EEG data of a similar experiment (Bartsch et al., 2015, experiment 3), we expected the GFBA effects of interest to be at least of medium effect size, for which G*Power (Faul et al., 2007) calculations would suggest twenty subjects to be sufficient for our within-subject repeated measures design (Cohen’s f > 0.25, power level 0.8, significance level 0.05).

### Stimuli and Procedure

#### Paradigm

We employed an unattended probe paradigm (UPP), which is a common experimental approach to investigate GFBA in humans and monkeys (Bartsch et al., 2018; C. J. McAdams & Maunsell, 2000; Saenz et al., 2002; Treue & Martínez Trujillo, 1999; Zhang & Luck, 2009). Participants are asked to covertly attend and discriminate a feature-defined target at a defined spatial location. Meanwhile, an unattended feature probe is presented elsewhere outside the current spatial focus of attention (FOA). Global feature attention will spread constantly across the whole visual field (Andersen et al., 2013; Liu & Mance, 2011), including both the location of the target and that of the distant unattended probe. The brain response to the probe is then analyzed as a function of the feature-similarity between the probe and the attended target. The response difference between a similar (matching) and a dissimilar (non-matching) feature-probe is taken as GFBA effect.

#### Stimuli

The stimuli are illustrated in Figure 2a. A half circle in the left visual field (VF) served as target and was presented together with a full circle in the right VF serving as a probe. Target and probe had a circle diameter of 3.1° visual angle and were placed 4.9° lateral and 3.1° below a central fixation cross. On each trial, the target was assigned one of two alternative colors (blockwise either red/green or blue/yellow) and the probe color was randomly chosen from a set of five colors (red, green, yellow, blue, magenta). The target represented either the left or right half of a circle, i.e., its convexity could either point to the left or right (changing randomly from trial-to-trial). Colors were psychophysically matched prior to the experiment in five independent participants via heterochromatic flicker photometry (Lee et al., 1988) with an average color luminance of 31 cd/m^2^. The background was dark grey (8.3 cd/m^2^), the fixation cross white (197 cd/m^2^).

#### Procedure

Participants were covertly attending to the target half circle in the left VF, while their fixation remained on the central cross. The participants were either asked to report the color of the target (red or blue: index finger, green or yellow: middle finger), or to report its orientation (convexity left: index finger, convexity right: middle finger). Task and target colors were designated at the beginning of the block. Every subject performed six color and six orientation blocks (each block lasting about 5min) in one of four possible pseudorandomized orders, such that the color of the targets (red/green or blue/yellow) was never repeated on subsequent blocks and the task (color or orientation) changed every second block. Target and probe were simultaneously presented for 300ms, followed by an inter-stimulus interval with only the fixation cross being present (randomly varying between 1000-1200ms, rectangular distribution). Each subject performed 180 trials per experimental block, yielding at least 216 trials for the individual trial types (averaged across different colors, see below).

#### Task types

In different experimental blocks, participants discriminated either the color (color task, attentional two-color template), or the orientation of the target (orientation task, color irrelevant). For both tasks, stimulus geometry and stimulus timing were kept identical, such that only the attention condition but no physical stimulation properties differed between task blocks. The purpose of the color task was to investigate the dynamics of biasing color selection for two colors ‘on the fly’ the moment target color A or B appears. The orientation task served as a control condition with color being irrelevant. That way, we could separate effects of attentional color biasing from effects of color selection that would also appear when color is irrelevant, e.g., caused by target discrimination itself, color priming (Kristjánsson, 2006; Theeuwes, 2013), or even low-level stimulus properties. In fact, any discrimination performed on the target will most likely entail some processing of its color as would be revealed by the orientation task.

#### Trial types and GFBA derivation

The probe’s color could match the color of the currently-presented target half circle (present target color, PC), match the distracting alternative target color (DC), or neither of them (non-target color). On a given trial block, there were always three non-target colors, i.e., the two colors that would be a target on the other blocks, and magenta which was added to introduce a greater variety of probe colors. However, since magenta was never used as a target, it is the only color where color-specific effects could not be controlled for (no attended vs. not attended comparison possible), such that magenta probes were excluded from further analyses. Figure 2b illustrates examples of the different trial types for a red probe. Effects of global feature-based attention (GFBA) were assessed by measuring the brain response to the color probe in the unattended hemifield (unattended probe paradigm, UPP). The brain response elicited by probes containing attended colors, i.e., the PC or DC were then compared with responses to probes drawn in unattended colors not relevant in the current experimental block (non-target colors, serving as baseline condition), see also previous work (Bartsch et al., 2015, 2017, 2018; Bondarenko et al., 2012). To increase the signal-to-noise ratio and to also avoid color-specific confounds, brain responses were averaged across different probe colors (red, green, blue, yellow) within trial-types. As visible in Figure 2b, the response to the same physical color probe could be compared under different attention conditions. However, the simplicity of the experimental design with a mono-colored target entailed limitations of color balance between trial types. Specifically, only when the probe matched the current target, the same color was present in both visual fields. Nevertheless, as shown in previous experiments, the feature bias for this trial type cannot be attributed to low level stimulus properties. Without attention to color, the mere presence of the same color on both sides of the visual field does not itself give rise to GFBA or GFBA-like modulations (experiment 4 of Bartsch et al. (2015), experiment 1 of Boehler et al. (2011), separate object condition).

### Data acquisition

Participants were equipped with a 32-electrode cap and seated in a dimmed, electrically and magnetically shielded recording booth (Vacuumschmelze, Hanau, Germany) in front of a partly transparent screen (COVILEX GmbH, Magdeburg, Germany) at a viewing distance of 1.0m. Stimuli were back-projected onto the screen using an LCD projector (DLA-G150CLE, JVC, Yokohama, Japan) placed outside the booth. Stimuli were delivered using Presentation® software (Neurobehavioral Systems Inc., Albany, CA). Participants gave responses with their index and middle finger of the right hand using a LUMItouch response system (Photon Control Inc., Burnaby, DC, Canada).

### EEG recording

The electroencephalogram (EEG) data were continuously recorded using a 32-electrode cap with mounted sintered Ag/AgCl electrodes (Easycap, Herrsching, Germany) and a Synamps amplifier system (NeuroScan, El Paso, TX). Electrode positions were chosen according to the international extended 10-20-system (American Electroencephalographic Society, 1994). Contact between electrodes and head surface was established using the abrasive electrolyte gel Abralyt light (Easycap, Herrsching, Germany), impedances were kept below 5kΩ at all electrode positions. An electrode at the right mastoid served as online reference during recording, data were then offline re-referenced to the weighted mean of the electric activity of the electrodes at the left and right mastoid according to (Luck, 2005, pp. 107–108). To track eye movements, electrodes were placed at the outer canthi of both eyes (bipolar derivation) and an electrode placed below the right eye (unipolar derivation) recording both the horizontal and vertical electrooculogram (EOG). EEG and EOG data were band-pass filtered (DC-50Hz) and digitized at a sampling frequency of 254.31 Hz. The magnetoencephalogram (MEG) was simultaneously recorded with a 248-sensor BTI Magnes 3600 whole-head magnetometer system (4D Neuroimaging, San Diego, CA, USA). Respective MEG data are not reported here.

### Data analysis

#### Behavioral data

Response time and response accuracy were computed using MATLAB routines (MathWorks Inc., Natick, MA, USA), respective temporal onsets and identities of stimuli and given responses were derived from the logfiles produced by Presentation® software. Trials with anticipatory (<200ms) and delayed (>1300ms) responses were excluded, and only correct responses were used for response time measures. For statistical validation, the data were analyzed with the software package SPSS (SPSS Inc., Chicago, IL, USA) by computing repeated measures ANOVAs (rANOVAs). Significant main or interaction effects were further evaluated using subsequent paired Student’s t-tests. An alpha of 0.05 served as significance level, Greenhouse-Geisser correction was applied to correct for non-sphericity.

#### EEG - Epoching and Artifact rejection

The continuous EEG data were epoched offline from 200ms before stimulus onset to 700ms after stimulus onset. After excluding anticipatory (<200ms), delayed (>1300ms), and incorrect responses, epochs were subjected to an artifact rejection. All epochs exceeding specific peak-to-peak amplitude measure thresholds were removed until the data were devoid of major artifacts including eye blinks, eye movements, and physiological noise like muscle tension. To this end, data were visually inspected and thresholds individually determined for every subject ranging from 70-115µV (mean 97.5µV) and leading to on average 5.4% rejected trials.

#### Event-related potentials (ERPs)

After artifact rejection, the remaining epochs (including only correct responses) were averaged locked to stimulus onset for the individual trial types within participants. Sensor sites and time ranges were chosen in accordance to previous work (Bartsch et al., 2015, 2017, 2018; Bondarenko et al., 2012). Specifically, GFBA effects are expected to appear between 150-300ms contralateral to the unattended probe at parieto-occipital sensor sites. More specifically, due to the contralateral retinotopic organization of the visual cortex, a probe in the right VF will elicit maximal GFBA activity in left ventral occipital cortex, that can typically be measured best at electrodes at parieto-occipital recording sites placed over the left hemisphere (i.e., PO3, PO7, PO9). A visual inspection of the data confirmed prominent GFBA modulations for color in the expected N1/N2 time range. As can be seen in the topographical EEG maps (Figure 4b, 7b, and 8b), effect maxima are fairly comparable to previous work with electrodes PO3 and PO7 appearing closest to topographic field maxima across all conditions. Hence, the signal was averaged across respective electrode sites for the reported ERP waveforms. ERP waveforms are plotted from −150 to 300ms with the 150ms pre-stimulus period serving as a baseline for all analyses. ERP waveforms and topographical field distribution maps were plotted using the Event-related Potential Software System ERPSS (Event-Related Potential Laboratory, University of California San Diego, La Jolla, CA, USA). A smoothing gaussian filter (low pass, half amplitude cutoff frequency of 23 Hz) was applied for visualization purposes only, statistical testing was performed on unfiltered data. All waveforms and topographical maps display ‘grand average’ data (i.e., data averaged across all twenty-two participants).

#### Statistical validation of amplitude differences

To retrieve the time course of GFBA modulations and define time windows of significant differences between the conditions, a time-sample by time-sample sliding 2×3 rANOVA was performed with the factors TASK (orientation, color) and COLOR (probe color matches PC, DC, or non-target). The data were tested in the time range of 0 to 300ms after stimulus onset, the width of the sliding window was 11.8ms (i.e., three time samples). To correct for multiple comparisons, we followed the logic of Wagner et al. (2017) taking into account the original sampling frequency (f_s_, here 254.31Hz) and the applied low-pass filter (f_c_, here 50Hz). The corrected alpha level was 1-(1-0.05)^2fc/fs^ ≈ 0.02. The first out of five or more successive sample points with a p-value below 0.02 was considered as effect onset. All subsequent statistical comparisons were performed within time ranges of significant main or interaction effects. For further explorative analyses, additional sliding t-tests between single conditions were performed outside the pre-defined time windows.

#### Median response time split analysis (RT split)

Data were re-analyzed to separate between slow and fast responses for specific trial types. To prevent any contamination by other influences like a particular color or response button (e.g., responses might be faster when the target is blue compared to green, or when answering with the index finger compared to the middle finger), the split was performed for every single stimulus display (each stimulus was shown 27 times, e.g., red target facing to the left with probe being green). Afterwards, all trials tagged as “slower” or “faster” than the median RT for that specific stimulus were averaged together with the respective “slow” or “fast” trials of the other stimuli contained in a specific trial type.

#### Target repetition analysis

Data were analyzed as a function of whether the target color was identical to that of the previous trial (target-repetition trials), or changed (target-switched trials). Since target color was randomly drawn from the two possible target colors on every trial, the analysis yielded equally-sized bins for the conditions (same probability for each trial to be a repeat or switch trial). Analogous to the RT split analysis, trial split was performed for every single stimulus display before averaging together the trials for the individual trial types (PC/DC/non-target).

### Data availability

All analyzed and generated datasets will be made freely accessible at https://osf.io/chbnd/.

## Acknowledgements

This work was supported by the Deutsche Forschungsgemeinschaft (Grant SFB779/TPA1). We thank Steffi Bachmann and Laura Hermann for assistance with the data acquisition.

## Additional Information

### Author contributions

MVB and JMH planned the experiments. MVB performed the research. MAS provided scientific support. MVB wrote the manuscript. JMH and CM edited and reviewed the manuscript.

